# Theoretical investigation of active listening behavior based on the echolocation of CF-FM bats

**DOI:** 10.1101/2021.12.23.474076

**Authors:** Takahiro Hiraga, Yasufumi Yamada, Ryo Kobayashi

**Affiliations:** Department of Mathematical and Life Sciences, Hiroshima University. C-201, Department of Sciences,1-3-2 Kagamiyama, Higashi-Hiroshima, Japan; Program of Mathematical and Life Sciences, Hiroshima University. C-623, Department of Sciences,1-3-2 Kagamiyama, Higashi-Hiroshima, Japan; Program of Mathematical and Life Sciences, Hiroshima University. C-201, Department of Sciences,1-3-2 Kagamiyama, Higashi-Hiroshima, Japan

**Keywords:** Three-dimensional spatial localization, Active listening behaviour, CF-FM bats, Interaural sound pressure level difference, High duty cycle echolocator, Mathematical analysis

## Abstract

Bats perceive the three-dimensional (3D) environment by emitting ultrasound pulses from their nose or mouth and receiving echoes through both ears. To detect the position of a target object, it is necessary to know the distance and direction of the target. Certain bat species synchronize the movement of their pinnae with pulse emission, and it is this behavior that enables 3D direction detection. However, the significance of bats’ ear motions remains unclear. In this study, we construct a model of an active listening system including the motion of the ears, and conduct mathematical investigations to clarify the importance of ear motion in 3D direction detection. The theory suggests that only certain ear motions, namely three-axis rotation, accomplish accurate and robust 3D direction detection. Our theoretical analysis also strongly supports the behavior whereby bats move their pinnae in the antiphase mode. In addition, we provide the conditions for ear motions to ensure accurate and robust direction detection, suggesting that simple shaped hearing directionality and well-selected uncomplicated ear motions are sufficient to achieve precise and robust 3D direction detection. Our findings and mathematical approach have the potential to be used in the design of active sensing systems in various engineering fields.

**Author Summary:** Many mammals use visual sensing for primary perception of their surroundings, whereas bats accomplish spatial perception by active acoustic sensing. In particular, by emitting ultrasound pulses and listening to the echoes, bats localize reflective objects, a process known as echolocation. Certain bat species move both of their ears while receiving the echoes, but the essential theory behind this ear movement remains unclear.

This paper describes a simple mathematical model for investigating the active listening strategy employed by bats. The theory suggests that the ear motions employed by bats enables highly accurate direction detection that is robust to observation errors. In addition, we determine what kind of ear motions are optimal for 3D direction detection. This study not only reveals the significance of pinnae motions in bats, but also opens up the possibility of engineering applications for active listening systems.

## Introduction

Bats perceive the three-dimensional (3D) environment by echolocation, which is the active ultrasound sensing capability to image their surroundings using the echoes reflected from surroundings by pulse emissions [1]. Despite the simple sensing design, equipped with only one transmitter (mouth or nose) and two receivers (left and right ears), bats accomplish precise navigation tasks in the air, such as the pursuit of prey [2, 3] and flying together with multiple conspecifics [4, 5]. The highly sophisticated mechanisms that enable 3D navigation with ultrasound have attracted extensive and longstanding attention from physiological and behavioral scientists.

To date, the acoustic imaging process in the auditory system has been widely investigated for bats and other animals [6-8]. Previous studies have reported that bats have an encoding mechanism for the Interaural sound pressure Level Difference (ILD) in the lateral superior olive, as seen in many mammals [8-13]. The lateral superior olive in bats has a relatively large capacity [14], and acoustic localization with ILD is physically suited to less diffractive high-frequency sound. Thus, the ILD encoding mechanism is regarded as a key property for 3D localization of bats. Recent studies have conducted more comprehensive analysis that combines ILD mechanisms with head-related transfer functions [15-17]. These functions are important features that describe the echo strength as a function of the echo arrival direction. Measurements of head-related transfer functions in various bat species suggest that the pinnae are used for beamforming to echoes reflected from objects[15-18].

This is not the only key evolutionary feature that bats have acquired for acoustic localization. Several species of bats employ behavioral solutions for 3D localization. *Rhinolophidae* and *Hipposideridae* families synchronize the movement of their left and right pinnae with pulse emission [19-22]. This active listening behavior has been reported for constant frequency–frequency modulated (CF-FM) bats, who use a compound signal consisting of a CF part and an FM part (Fig 1A). Previous physiological and ethological studies have clarified that CF-FM bats detect the precise time interval between pulse emission and echo arrival using the FM part to measure the distance to the object accurately [7, 23]. The CF part is used for fluttering moth detection and Doppler shift compensation [24-26]. According to measurements from *Rhinolophus ferrumequinum*, both pinnae move continuously while listening to the CF part of the echo [20]. Based on this behavioral evidence, several studies have investigated the usefulness of ear motions for 3D localization through mathematical simulations [27, 28] or practical demonstrations [20, 29], but the essence of appropriate pinnae motions is still unclear.

**Fig 1.**
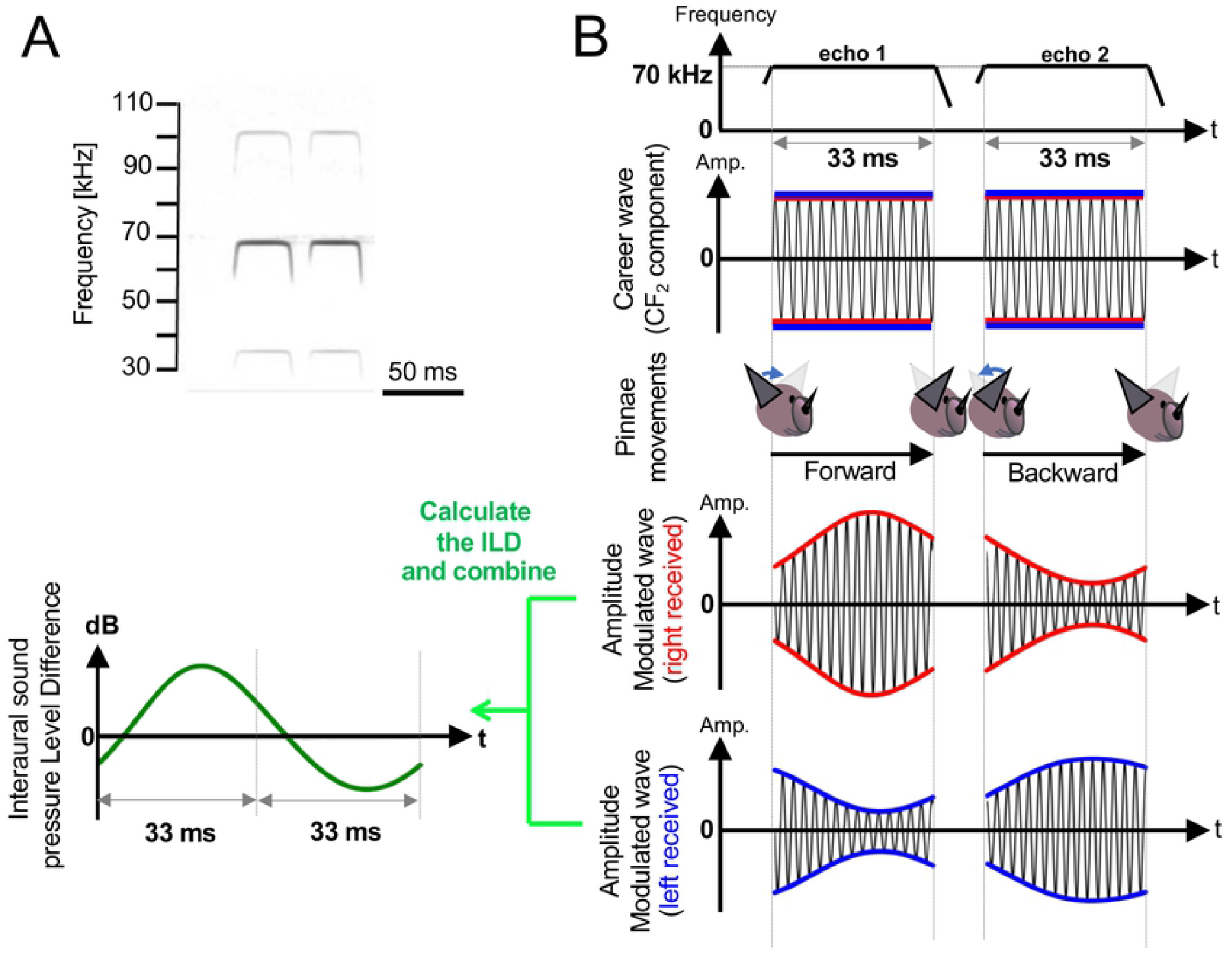
Pulse emission and reflective echo patterns. (A) Typical time-frequency structure of the echolocation pulse emitted by *Rhinolophus ferrumequinum nippon*. (B) Schematic diagram of amplitude modulation of CF_2_ component in the echo caused by pinnae motions.

Even if precise 3D pinnae motions could be measured, it would be difficult to determine their exact effects because bat behavior always exhibits the best-benefit response. In contrast, a theoretical approach allows us to evaluate various pinnae motions, including those of bats. Moreover, theoretical investigation can isolate the various factors of acoustic localization and provide insights into their essential components, give an interpretation of bat behavior, and possibly provide support for biomimetic applications.

Based on these motivations, exhaustive ear motions were analyzed to identify the underlying theory of appropriate ear motions. In these analyses, various ear motions were evaluated in terms of their 3D direction detection performance using custom-made functions and supervised machine learning.

## Methods

### Behavioral traits of bats reflected in our model

In this subsection, we describe the behavioral traits of bats reflected in our model. Fig 1A shows a typical time-frequency structure of the echolocation pulses emitted by CF-FM bats (*Rhinolophus ferrumequinum nippon*) recorded in previous study[30]. In these pulses, the energy maximum appears in the second harmonic of the CF part (CF_2_); bats actively use CF_2_ for fluttering moth detection and Doppler shift compensation [24-26]. To simplify our simulations, amplitude modulation was only calculated for the CF_2_ component of the echo.

According to previous studies that measured the ear motions of bats, *Rhinolophus ferrumequinum* continuously move their pinnae while listening to the CF part of the echo [20]. These bats adjust their left and right pinnae in an antiphase manner [19, 20]. In particular, the pitch angle of the ears tends to move from back to front or from front to back while listening to the echoes, which can be modeled as a cosine phase [19]. Based on these findings, asymmetrical ear motions were embedded in bat mimicking simulations.

Fig 1B shows a schematic diagram of the amplitude modulation of CF_2_ in the echo caused by pinnae motions. Because CF-FM bats tend to conduct the sensing process twice in the space of one periodic pinnae motion [19], echo signals obtained from two sensing operations were simulated in our analyses. With reference to previous measurements of *Rhinolophus ferrumequinum* [20], the echo frequency was set to 70 kHz (i.e., wavelength *λ* = 5 mm) and the echo duration was set to 33 ms. Note that silence time between 1^st^ and 2^nd^ echo was removed, and both signals were combined.

### Model of the direction detection system

Fig 2A shows a schematic diagram of the environmental setup for the left and right ears and a target object. A single target object was stationed in the direction expressed by the azimuth angle *θ* and elevation angle *φ*, or equivalently by the unit vector ***n*** = (cos *θ* cos *φ*, sin *θ* cos *φ*, sin *φ*), which we call the direction vector. In our model, the amplitude modulation of the echo is caused by changes in the directional attitude of the ears. Fig 2B shows a schematic diagram of the left and right ears and a speaker when all materials are directed in front of the bat (positive direction of x-axis). To construct a directional ear, four omni-directional microphones were placed at the vertices of a rectangle. The four echo signals obtained from these microphones were summed to generate the overall received signal. In particular, by adjusting the horizontal and vertical spacing between the microphones (*δ*_*y*_, *δ*_*z*_), the hearing directivity pattern could be controlled. Fig 2C shows the hearing directivity pattern used in this study. Based on measurements and computational representations of the hearing directivity patterns of CF-FM bats, including *Pteronotus parnellii* [17, 18], *Hipposideros pratti* [31], *Rhinolophus Roxi* [17], and *Rhinolophus ferrumequinum* [31], the half-amplitude angle (−6 dB off-axis angle from the maximum sensitivity angle) tends to be distributed from 40–90° off the horizontal axis. In addition, the directivity forms an asymmetrical 3D pattern. Based on these characteristics, *δ*_*y*_, *δ*_*z*_ were set to be slightly smaller than half of the echo wavelength λ. As a result, an asymmetrical beampattern was reproduced, as shown in Fig 2C.

**Fig 2.**
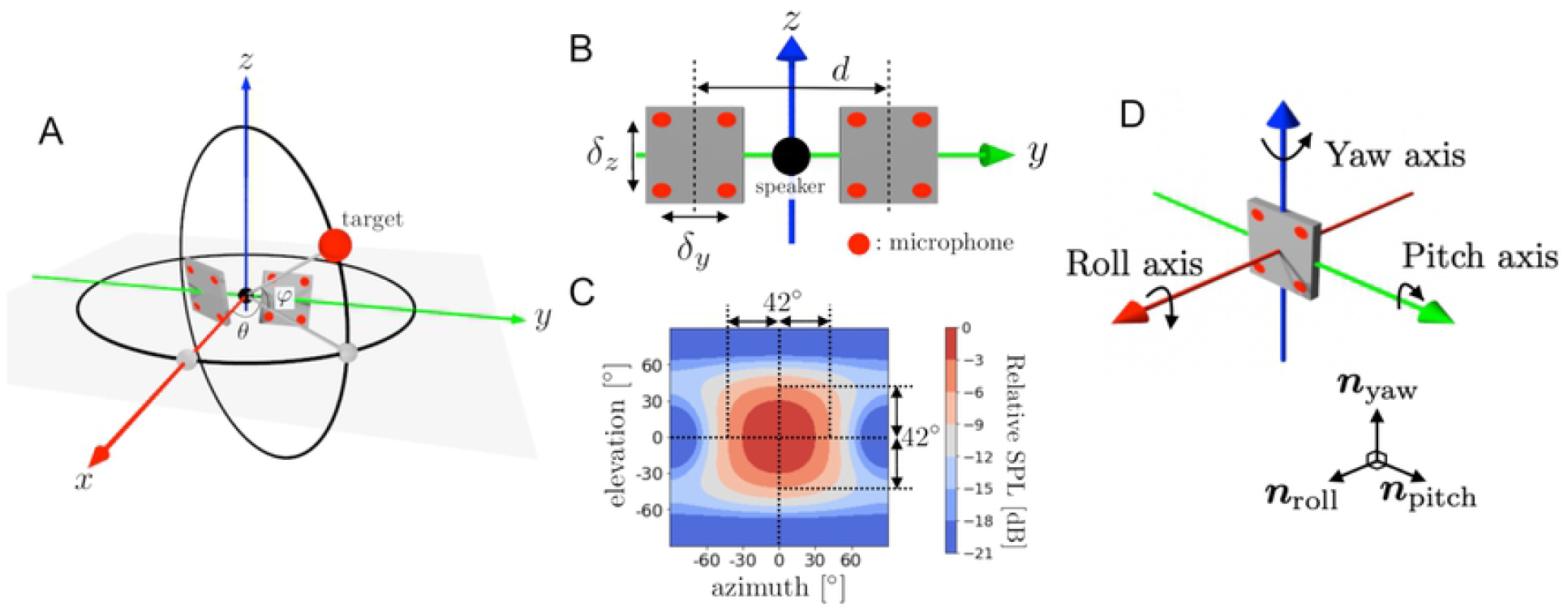
Schematic diagram of model setup. (A) Direction of the target expressed by azimuth angle *θ* and elevation angle *φ*. (B) Positions of the two directional ears with spacing *d*. Each ear consists of four omni-directional microphones, where *δ*_*y*_ and *δ*_*z*_ are the horizontal and vertical spacings of each microphone. (C) Hearing directivity pattern of the ear. (D) Three axes (roll, pitch, yaw) fixed to the directional ear and corresponding orthonormal basis [***n***_*roll*_,***n***_*pitch*_, ***n***_*yaw*_].

As shown in Fig 2D, the roll axis, pitch axis, and yaw axis are fixed to the directional ear and the unit vectors ***n***_*roll*_,***n***_*pitch*_, ***n***_*yaw*_ indicate the directions of these three axes. The attitude of the directional ear is then given by the matrix *L* = [***n***_*roll*_,***n***_*pitch*_,***n***_*yaw*_] ∈ *SO*(3). Additionally, the attitude change caused by the motion of the directional ear is expressed by the *SO*(3)-valued function *L*(*t*) = [***n***_*roll*_(*t*),***n***_*pitch*_(*t*),***n***_*yaw*_(*t*)], where *t* is the time variable. Assume that the target object is pointed to by the direction vector ***n*** and the echo received at the origin is a sinusoidal wave with amplitude *A* and wavelength λ. The directional ear in proximity to the origin receives a signal whose envelope component *S*_*env*_ is expressed by the following formula (see **S1 text**):

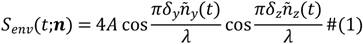

where *L*(*t*)^*T*^***n*** = ***ñ***(*t*)=(*ñ*_*x*_(*t*), *ñ*_*y*_(*t*), *ñ*_*z*_(*t*)). Therefore, the amplitude modulation of the echo envelope caused by the motion of the directional ear can be calculated for every target direction ***n*** once the attitude history *L*(*t*) is known. Note that *L*(*t*) and *S*_*env*_ (*t;****n***) are to be defined for the left and right directional ears. From the envelope of the left and right received echoes, the ILD is defined by the following equation:

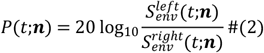

where ***n*** is the direction vector to the target and 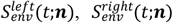 indicate the envelope of the left and the right received echoes under attitude histories *L*^*left*^(*t*) and *L*^*right*^(*t*), respectively.

The procedure described above obtains the ILD, which is a temporal signal *P*(*t;****n***), from the direction vector ***n***. Our question is whether we can obtain the direction vector ***n*** from the ILD signal *P*(*t;****n***). If so, what motions of the left and right directional ears make it possible, and how robust is the detection performance to observation errors?

### Evaluation function and degree of injection

To evaluate the effectiveness of the left and right motions of the directional ears mathematically, we introduce a general evaluation function and an index which we call the *degree of injection*. Let *X* be a set of state variables of the objective system, which we are going to identify through the observations. We write the observation process as the map

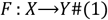

where *Y* is the space in which the observed data lie (possibly a Euclidian space or a functional space). Of course, we can define the map *F* only when the states of the system having the same state variable of *X* give the same observation data; hereafter, this is assumed to be true. To determine the state variable uniquely from the observed data, we require the inverse map

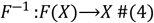

Therefore, the observation map *F* should be *injective*. In addition, to be sufficiently robust to observation errors, *F* must be non-degenerate, and hopefully not nearly degenerate at any point in *X*. (Here, ‘degenerate’ means that the dimension of the tangential map’s image is less than the dimension of *X*.) Based on these considerations, we define the evaluation function *U*_*F*_ on *X* as follows:

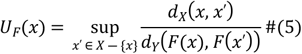

where *d*_*X*_ and *d*_*Y*_ indicate the distance functions defined in spaces *X* and *Y*, respectively. *U*_*F*_(*x*) = +∞ holds if the injective property of *F* is violated at *x* (meaning the existence of *x*′ ≠ *x* satisfying *F*(*x*′) = *F*(*x*)). In addition, *U*_*F*_(*x*) can measure the degree of degeneration of *F* at *x*. Actually, *U*_*F*_(*x*) becomes infinite if *F* is degenerate at *x*, and it attains a large value if *F* is nearly degenerate at *x*, which means that the inverse map is too sensitive to observation error at *F*(*x*). In any case, the large magnitude of the evaluation function *U*_*F*_(*x*) implies difficulty in constructing an inverse map or a well-behaved inverse map at *F*(*x*).

Finally, we define the degree of injection of *F* by the following equation:

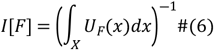

Note that *X* is usually a subset of some Euclidian space, and so the integral is definable. Large values of *I*[*F*] indicate that the evaluation function *U*_*F*_ does not take a large value in the state variable space *X*, so the well-behaved inverse map *F*^−1^ is expected to exist globally. This implies that the observed data contain rich information for determining the desired state variable. Conversely, if *I*[*F*] is small, *F*^−1^ itself or a well-behaved *F*^−1^ is difficult to construct.

Our task is to find the direction of the target from the time series data of the ILD. Thus, we consider *X* as a set of directions expressed by some subset of the unit sphere *S*^2^, for example,

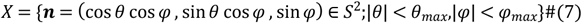

with the 2-norm in ℝ^3^. We set the measured data space to *Y* = *C*^0^([0, *T*]) with the sup-norm, where *T* is the period of the ear motions. In our problem, the observation process is determined by the attitude change of the left and right directional ears, expressed by the two *SO*(3)-valued functions *L*^*left*^(*t*) and *L*^*right*^(*t*) with period *T*. We denote the pair *L*^*left*^(*t*) and *L*^*right*^(*t*) as *M*, and use the notation *P*_*M*_(*t* ;***n***) for the ILD signal obtained by the attitude change *M* = (*L*^*left*^(*t*), *L*^*right*^(*t*)). We adopt the same symbol *M* for the map *M* : *X*→*Y* defined by

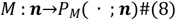

Note that the map *M* is definable because *P*_*M*_(· ; ***n***) is a function of the ratio between the amplitudes of the left/right envelope signals, which does not depend on the target distance and other factors like the reflection rate of the object. Following expression (5), we write the evaluation function as

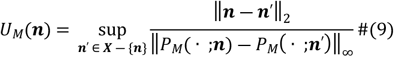

and we define the degree of injection of *M* by

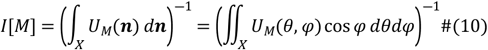

Using this index, we will evaluate various types of ear motions and compare them with the quality of the inverse map (pseudo-inverse map in the case of non-injectivity, as discussed later) constructed by the neural network described in the next section. Note that the expression (*θ,φ*) will often be used instead of the direction vector ***n***, as seen in (10), where this will not cause confusion.

### Evaluation of localization performance by supervised machine learning

Supervised machine learning is a good tool for constructing an inverse map numerically when an analytical expression is intractable. To confirm that the inverse map can be constructed when the appropriate ear motions are employed, a 3D direction detection test was conducted using a fully connected neural network. Fig 3 shows a schematic diagram of a fully connected neural network and the data flow. Supervised machine learning was performed using this network. The input data to the neural net were the discretized ILD data calculated from the angle pair (*θ, φ*) under the adopted ear motion *M*, and the output data were the angle pair (*θ*_*guess*_, *φ*_*guess*_), i.e., the estimated (*θ, φ*). The detection error in constructing inverse map was evaluated by the following equation,

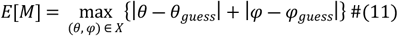

**Fig 3.**
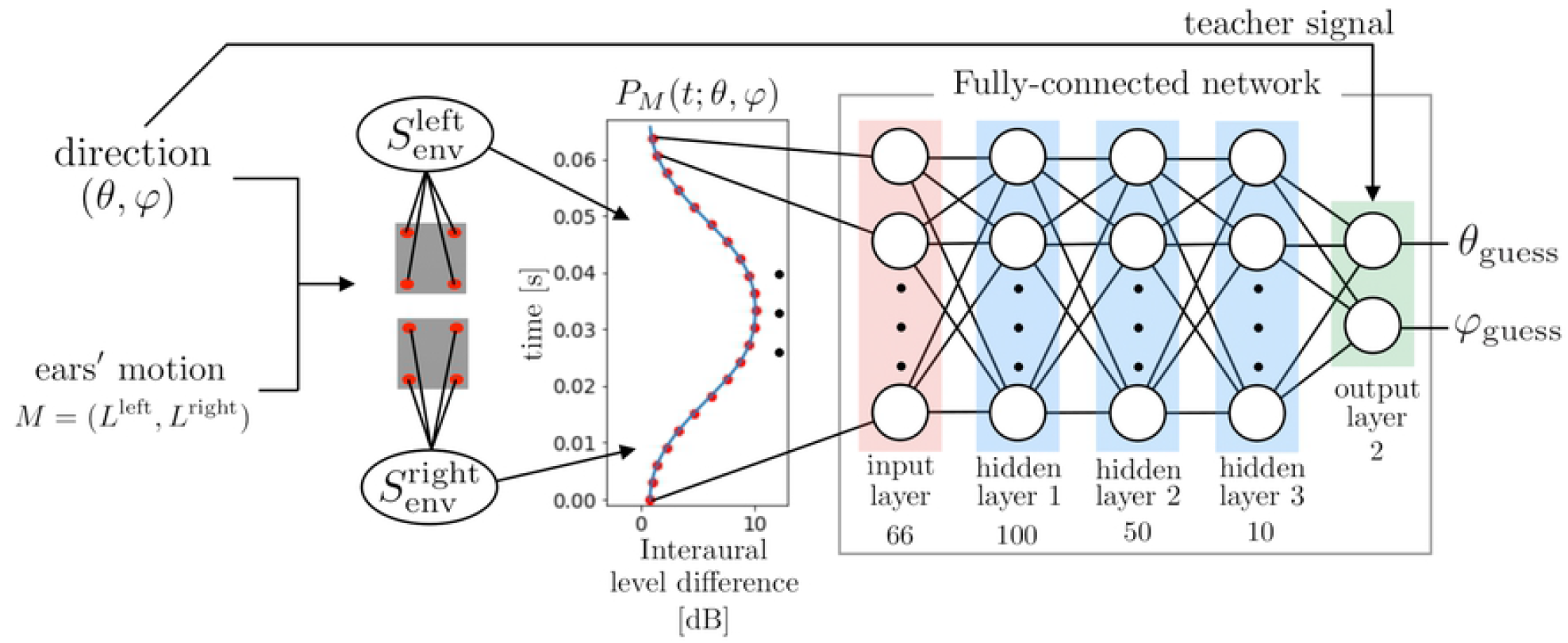
Schematic diagram of the supervised learning approach for obtaining the inverse map of *M*. The ILD signal is calculated for all directions (*θ, φ*)∈*X* fixing the ear motions. It is discretized at time intervals of 1 ms and passed to the input layer of the neural network. In the neural network, the ReLU activation function is used in hidden layers 1, 2, and 3, and the mean squared error is the error function in the output layer.

The azimuth angle *θ* and the elevation angle *φ* were restricted within ±60°. The neural network was trained 5000 times using uniformly distributed random (*θ, φ*) data. During the last 250 steps of the training, tests were carried out between every training step. In the test condition, the azimuth angle *θ* and the elevation angle *φ* were divided into 5.45° increments so that 23×23 situations were tested, and the detection errors were evaluated for every tested angle pair (*θ, φ*). Finally, *θ*_*guess*_ and *φ*_*guess*_ are evaluated as the median of the last 250 output data, respectively.

NOTE: Under the supervised learning approach described above, the inverse map of *M* is constructed when *M* is injective, in some accuracy level. However, our network learns some inverse-like map even when *M* is not injective, which we call the pseudo-inverse map. This pseudo-inverse map works as follows:

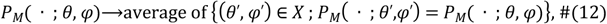

where ‘average’ means the center of gravity in the *θ*-*φ* plane in this case.

### Setting of directional ear motion patterns

The specific form of the directional ear motions can be written as follows using the roll–pitch–yaw expression (see **S2 Text**):

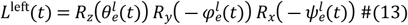

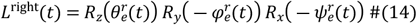

where the six angle functions 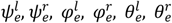 are periodic with period *T*, and the frequency of the ear motions is set to *f*_*e*_ = *T*^−1^. In our model, the periodic motion is restricted to the 0^th^ and 1^st^ Fourier modes, because actual bats do not exhibit complicated motion [19, 20]. Thus, we define the pairing types of the left- and right-ear angle functions as listed in Table 1. In our simulations, the roll, pitch, and yaw angle functions (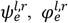 and 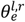) were chosen from the pairing types listed in Table 1.

**Table 1.** Pairing types of left and right angle functions 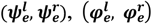, and 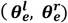.

| Pairing name | Left angle function | Right angle function |
| --- | --- | --- |
| **0** | 0 | 0 |
| **$\overline{\text{CONST}}$** | $C$ | $-C$ |
| **SIN** | $C \sin(2\pi f_e t)$ | $C \sin(2\pi f_e t)$ |
| **$\overline{\text{SIN}}$** | $C \sin(2\pi f_e t)$ | $-C \sin(2\pi f_e t)$ |
| **COS** | $C \cos(2\pi f_e t)$ | $C \cos(2\pi f_e t)$ |
| **$\overline{\text{COS}}$** | $C \cos(2\pi f_e t)$ | $-C \cos(2\pi f_e t)$ |

## Results

### Typical examples for direction detection with ear motions

To confirm the usefulness of the ear motions, two patterns (with and without ear motions) were compared. Fig 4 shows the evaluation function *U*_*M*_(*θ, φ*) and the results of machine learning under two patterns: 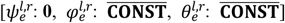 as a static example and 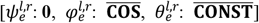 as a dynamic example. As shown in Figs 4 A2-3 and B2-3, the colormap of *U*_*M*_(*θ, φ*) reflects the geometric pattern of the distribution of detection errors by the neural network. The degree of injection *I*[*M*] is less than 0.001 for the static condition and 0.24 for the dynamic condition. Moreover, the detection error *E*[*M*] is 109.4° for the static condition and 16.9° for the dynamic condition.

**Fig 4.**
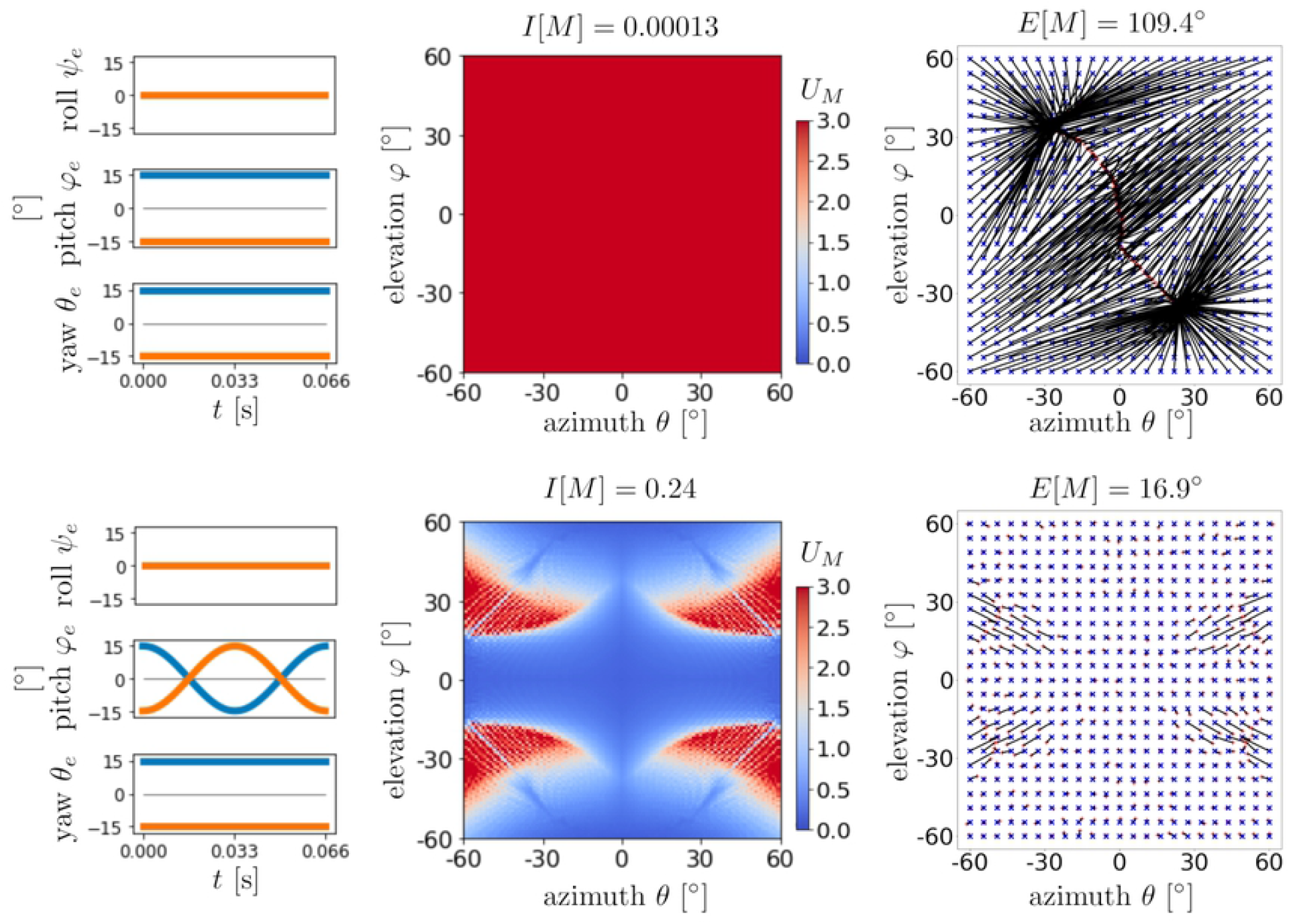
Examples of direction detection performance with and without ear motions. (A1, B1): Combination of angle functions. (A2, B2): Colormaps of evaluation function *U*_*M*_(*θ,φ*) and the degree of injection *I*[*M*]. (A3, B3): Results of machine learning. Blue ‘x’ markers indicate test data (*θ,φ*) and red ‘+’ markers indicate output data (*θ*_*guess*,_ *φ*_*guess*_). Black lines are the error lines connecting points (*θ,φ*) and (*θ*_*guess*,_ *φ*_*guess*_). The detection error *E*[*M*] is also given.

Examples of more complete direction detection are shown in Fig 5. In these examples, the ear motions conditions were chosen as 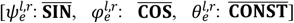 and 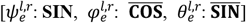. In each condition, the evaluation function *U*_*M*_(*θ, φ*) takes smaller values in the whole domain, and the degrees of injection *I*[*M*] are 1.52 and 1.35, respectively. The detection errors *E*[*M*] are 2.1° and 2.5°, indicating that accurate direction detection is accomplished. These results suggest that it is necessary to combine the roll, pitch, and yaw rotations appropriately for accurate detection of the 3D direction. Additionally, the results in Figs 4 and 5 indicate that the degree of injection is strongly related to the direction detection performance.

**Fig 5.**
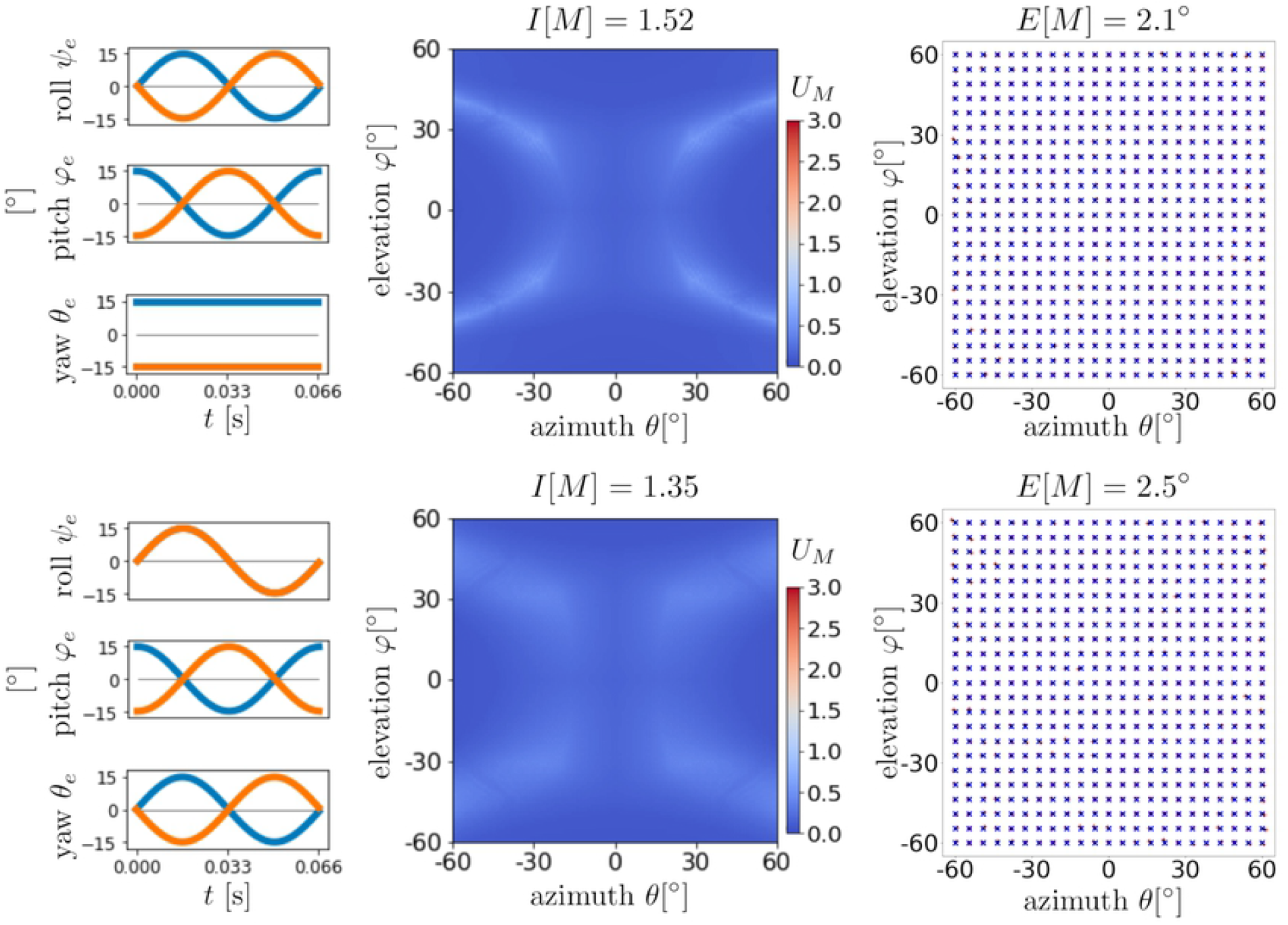
Examples of direction detection performance with appropriate ear motions. The formation of Fig 5 is same as Fig4. Blue color map and less-visible error lines mean the good performance of direction detection.

### Exhaustive analysis of ear motions in pitch anti-phase case

To determine appropriate combinations of the roll, pitch, and yaw rotations, 36 motion patterns were analyzed. The corresponding evaluation functions and degrees of injection are shown in Fig 6. As described before, based on the actual motions of bats’ pinnae, the pitch angle functions 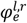 are fixed to the anti-phase pairing pattern 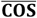.

**Fig 6.**
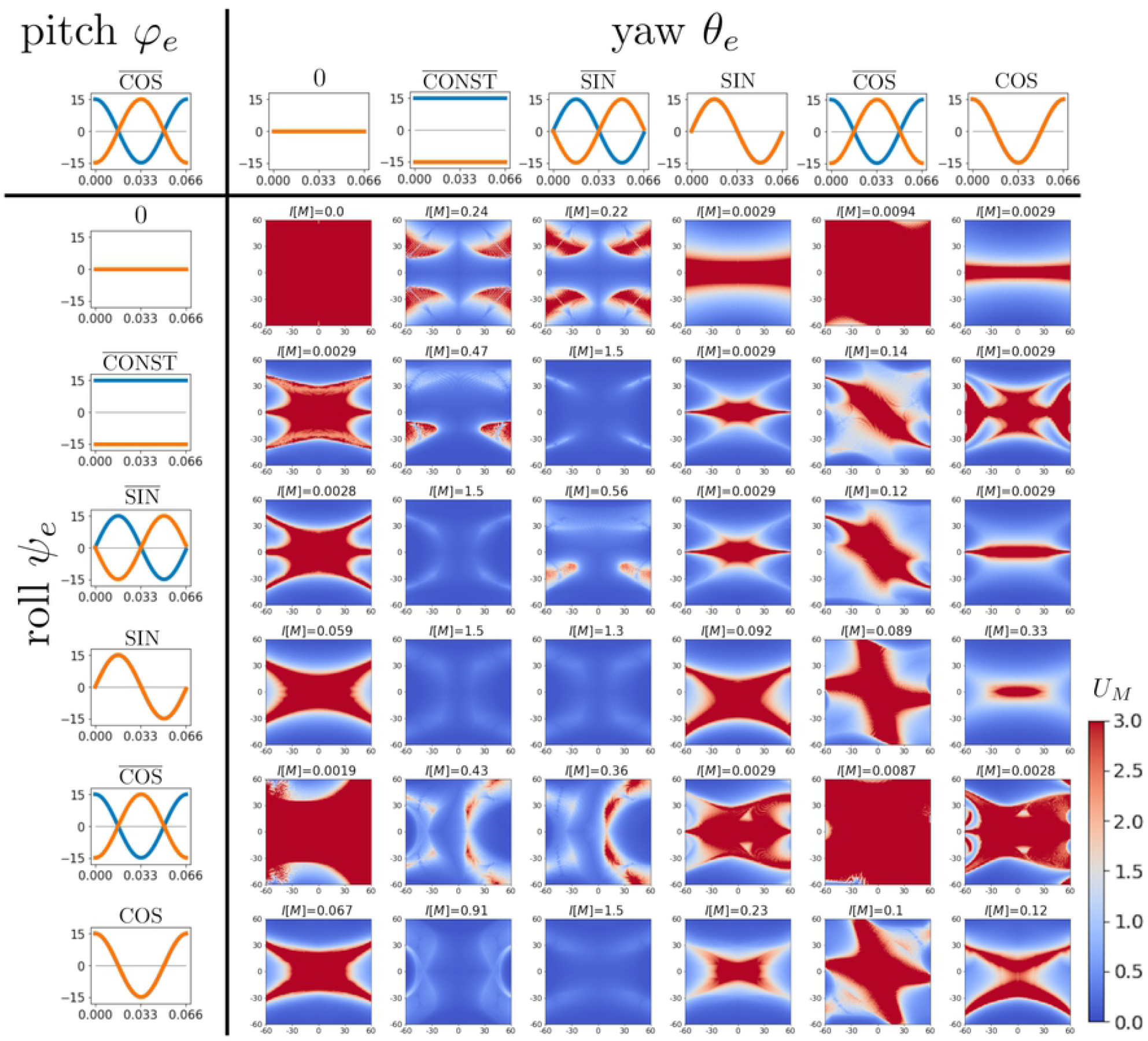
Colormaps of *U*_*M*_(*θ,φ*) and the degrees of injection for various ear motion patterns. The pitch angle functions 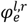 are fixed to 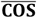 according to actual bat behavior. Blue and orange lines indicate the angle functions of the left and right ears, respectively. The left and top array panels display the roll angle functions 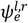 and the yaw angle functions 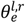, respectively.

To classify the ear motion patterns graphically, we focus on the orbits of ear motions given by 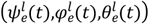 and 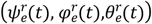 in *ψ*_*e*_ − *φ*_*e*_ − *θ*_*e*_ space. Additionally, the convex hull of the union of the left and right ears’ orbits in *ψ*_*e*_ − *φ*_*e*_ − *θ*_*e*_ space is considered. We classify the motion patterns according to the pair of dimensions of the convex hull and each ear’s orbit. As shown in Fig 7, there are five types of dimension pairs: 3-2, 3-1, 2-2, 2-1, and 1-1.

**Fig 7.**
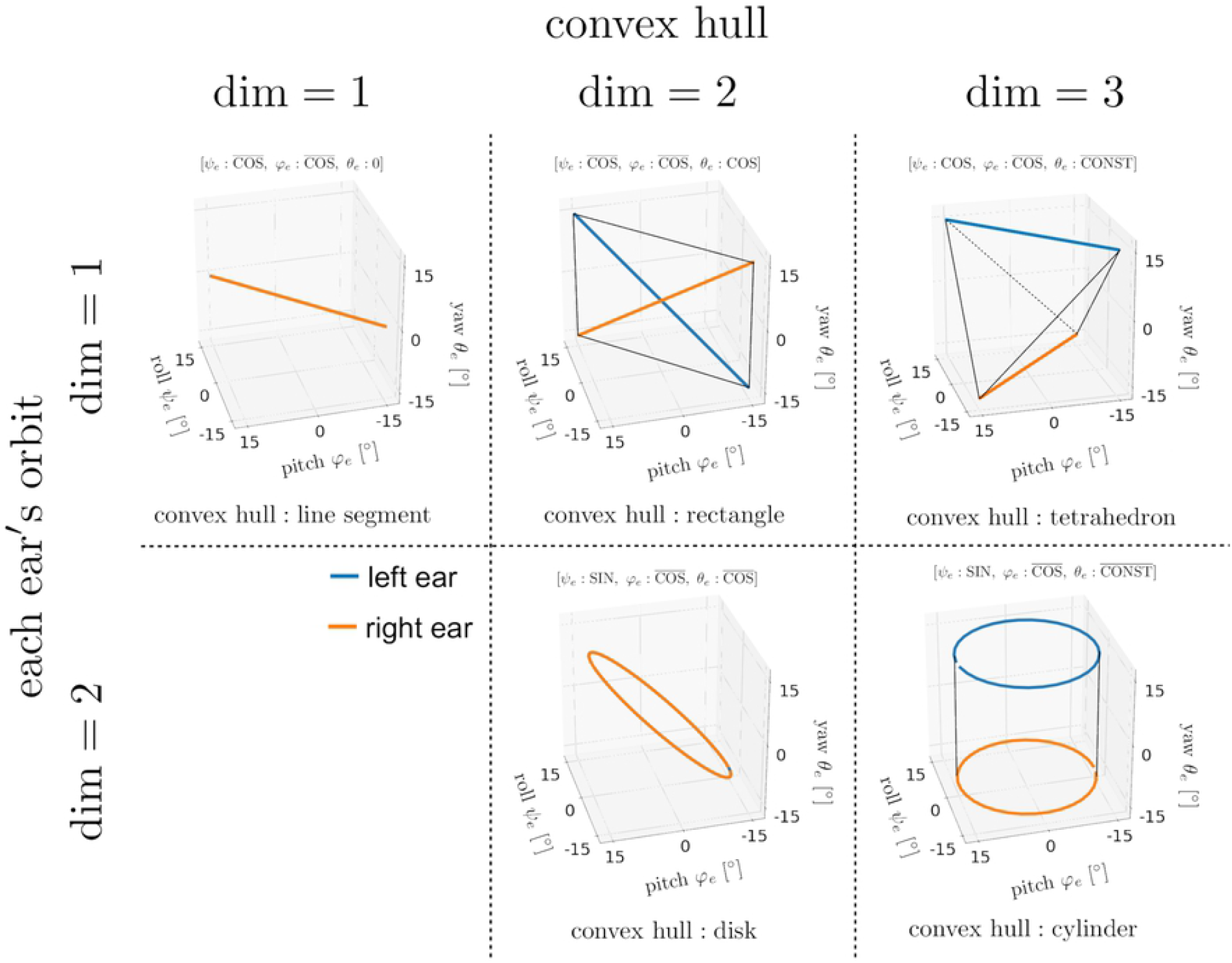
Five types of dimension pairs of the convex hull and each ear’s orbit. The blue lines indicate the left ear’s orbit 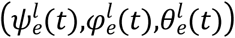 and the orange lines indicate the right ear’s orbit 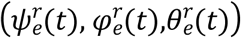. When both orbits coincide, only the orange line is displayed. The convex hull of the union of both ears’ orbits is displayed in each case.

Fig 8 exhibits the dimension pairs of the convex hull and each ear’s orbit, the degree of injection *I*[*M*], and the detection errors *E*[*M*] of the 36 motion patterns. There are 12 motion patterns (colored boxes) that achieve precise direction detection. Among them, 5 motion patterns (boxes bounded by red lines) have larger injection degrees, essentially indicating good motion patterns, as shown in the next subsection.

**Fig 8.**
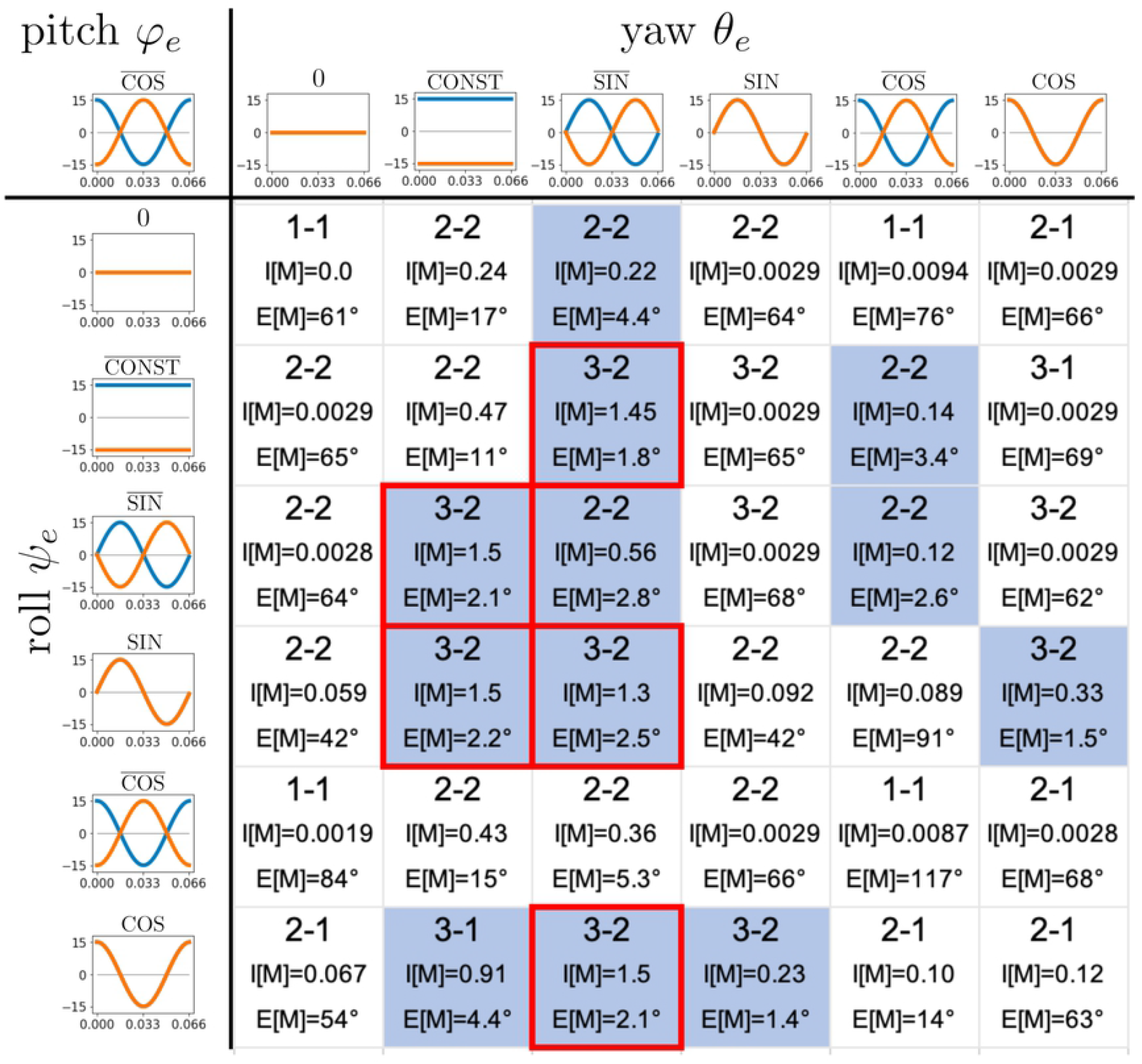
Dimension pairs and direction detection errors for various motion patterns. In each box, the dimension pair of the convex hull and each ear’s orbit is given in the upper part, the degree of injection is given in the middle, and the detection error is given at the bottom. Here, we adopt *E*[*M*] < 5° as the criterion for precise direction detection. The colored boxes indicate that the corresponding motion patterns give precise direction detection. The boxes bounded by red lines correspond to the motion patterns with large degrees of injection (*I*[*M*] > 1).

### Robustness against degradation of the ILD resolution

Next, the robustness of direction detection against the degradation of the ILD resolution was investigated. Fig 9A shows the relationship between the degree of injection and the direction detection error for the 36 ear motion patterns without the degradation of the ILD resolution (see green line in Fig 9B). We examined the detection robustness against the degradation of the ILD resolution for the relatively small detection error group (i.e., *E*[*M*] < 20°). The detection errors were reevaluated by decreasing the ILD resolution to 1 dB and 3 dB (see the orange and blue lines in Fig 9B). As shown in Fig 9C, the detection errors remained small for the group with larger degrees of injection (*I*[*M*] > 1), while the errors increased much more in the other groups. These findings suggest that 5 motion patterns satisfying conditions *I*[*M*] > 1 not only accomplish accurate direction detection, but are also robust to the degradation of the ILD resolution. From these characteristics and Fig 8, we can identify three ear motion conditions that ensure the precise and robust direction detection:

i. The convex hull of the union of the two ear orbits is three-dimensional;
ii. Neither orbit degenerates to one dimension;
iii. The left and right yaw angle functions do not coincide.

**Fig 9.**
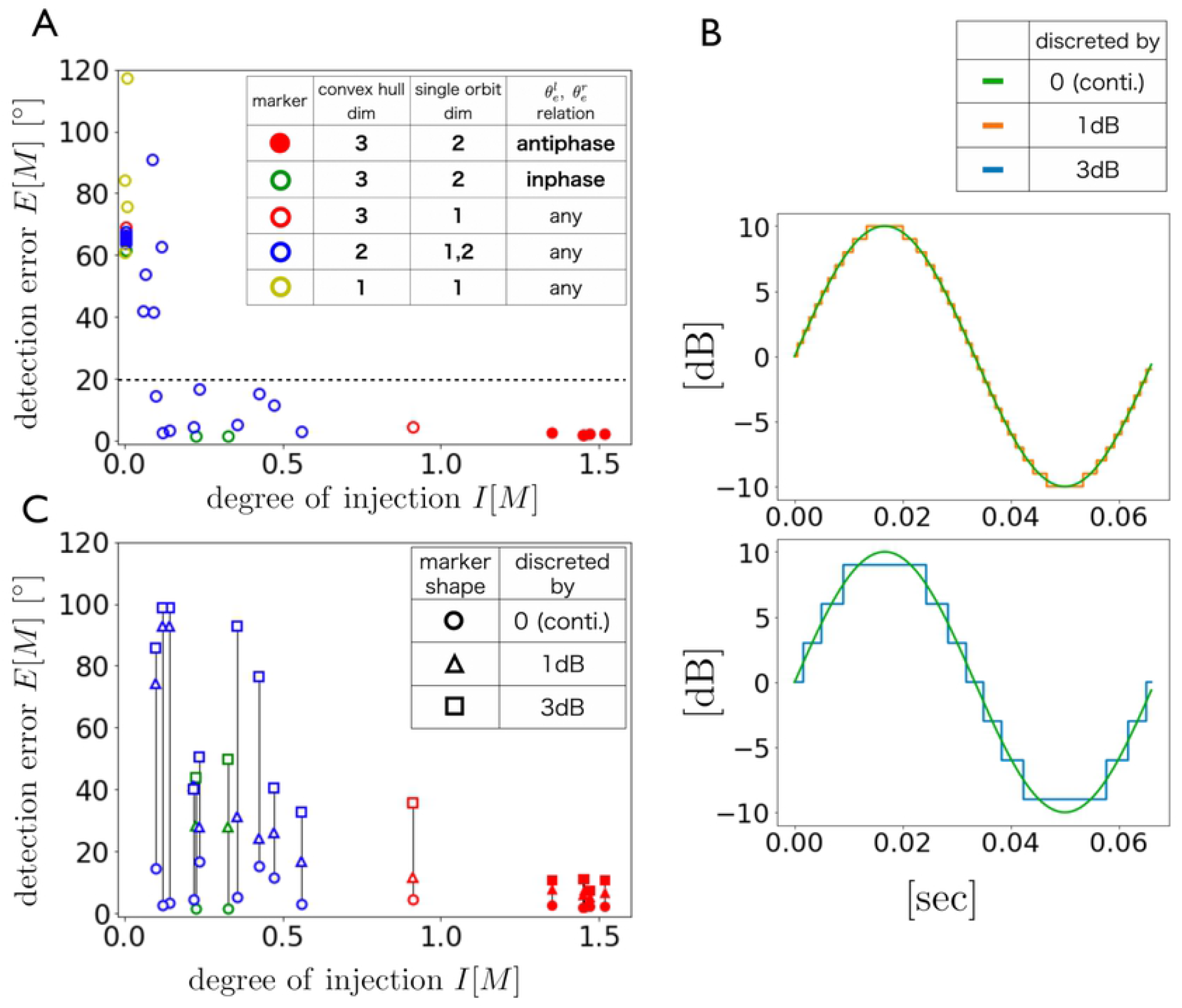
Relationship between *I*[*M*] and *E*[*M*] under various degradation levels of ILD resolution. (A) Relationship between the degree of injection and the detection error for the 36 ear motions without the degradation of the ILD resolution. (B) Example of change in the sinusoidal signal for each degradation level. (C) Relationship between the degree of injection and the detection error for each ear motion under the degraded ILD resolutions. Note that these evaluations were conducted for ear motions with relatively small detection errors (*E*[*M*] < 20°) in the no degradation condition (A). The length of the vertical black line corresponds to the increase in the detection error when the ILD discretization level changes from 0 dB to 3 dB.

### General case analysis

We now examine the general case. By removing the bat-motivated limitation of pitch motion 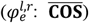, the detection performances were evaluated for 6^3^ = 216 ear motions in terms of the degree of injection *I*[*M*], as shown in Fig 10. These analyses show that the degree of injection *I*[*M*] is small when the angle relations 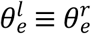 OR 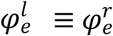 hold. Through these analyses, we determined the following conditions for ear motions satisfying *I*[*M*] > 1:

i. The convex hull of the union of the two ear orbits is three-dimensional;
ii. Neither orbit degenerates to one dimension;
iii. The left and right yaw angle functions do not coincide;
iv. The left and right pitch angle functions do not coincide.

**Fig 10.**
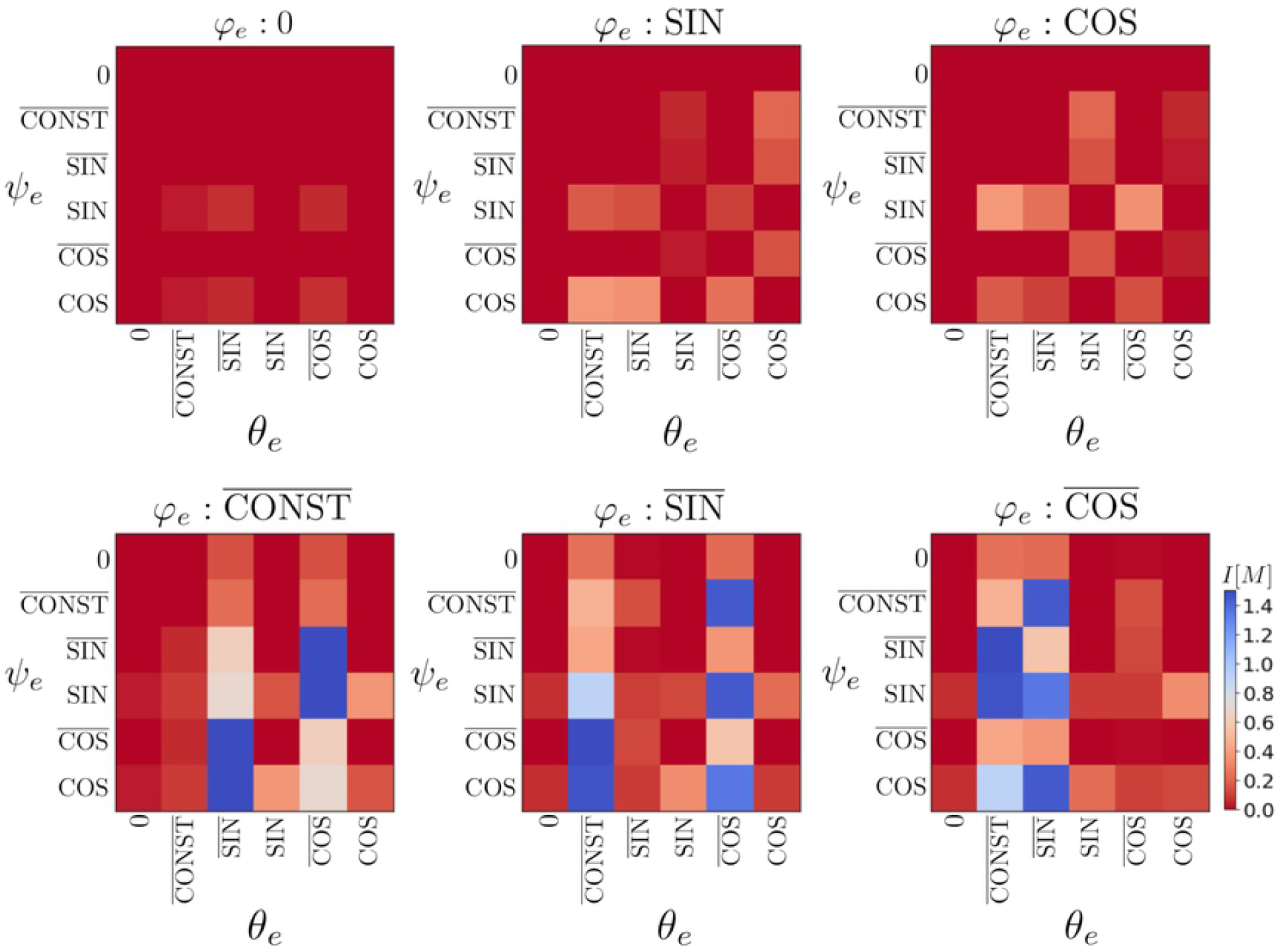
Colormaps of degree of injection *I*[*M*] of all combinations of *ψ*_*e*_ -*φ*_*e*_ - *θ*_*e*_ angle functions. The fixation of the pitch angle functions 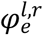 to 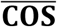 is removed, so that the degrees of injection were evaluated for 6^3^ = 216 motion patterns.

The 14 of 216 motion patterns satisfy the above four conditions. We confirmed that these 14 motion patterns achieve the precise and robust direction detection, and the other patterns do not.

Finally, the effect of phase differences in the left and right ear motions on the detection performance is examined in Fig 11. All motions have the same orbits, but the simultaneous lines vary according to the pitch–yaw (*φ*_*e*_-*θ*_*e*_) phase difference. This result suggests that phase differences larger than several tens of degrees is sufficient to achieve good-quality detection.

**Fig 11.**
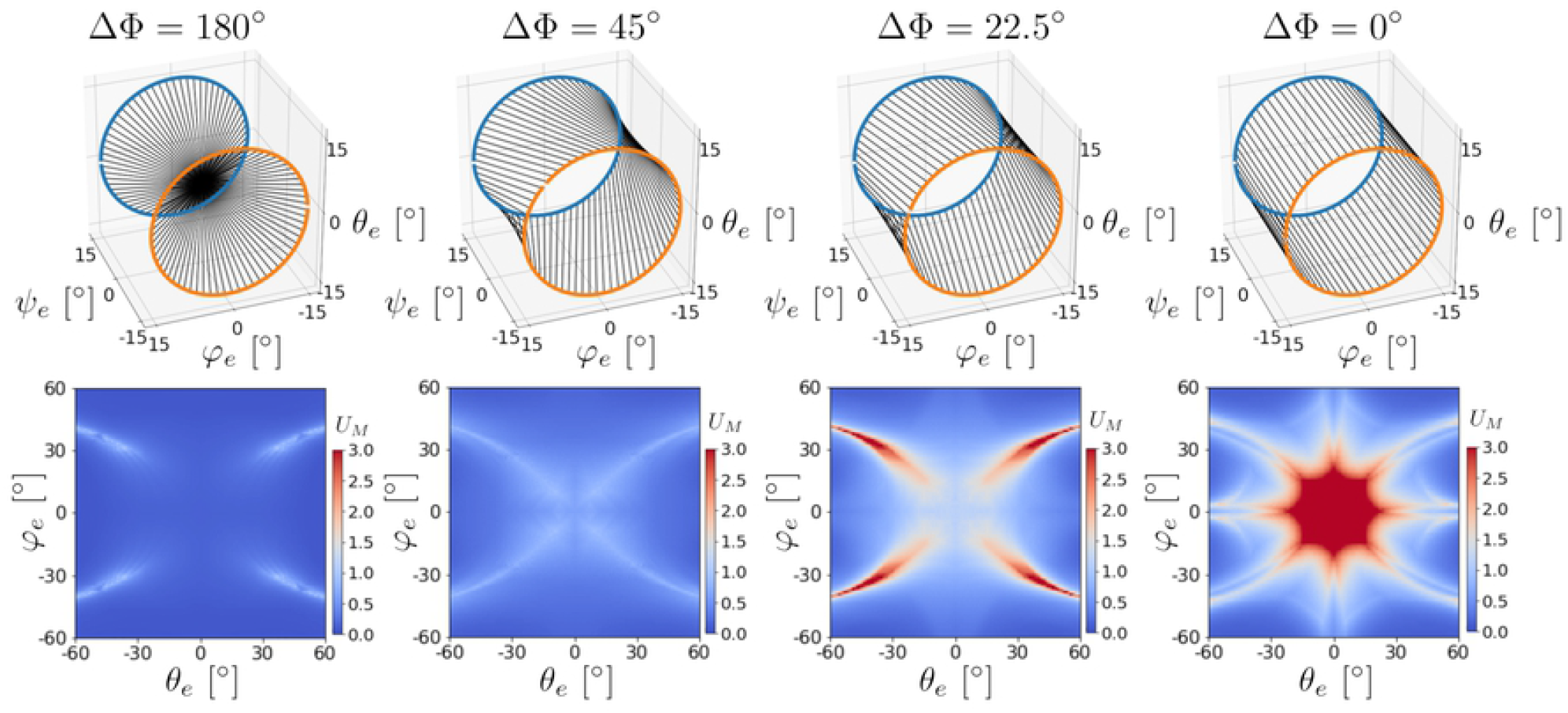
Effect of phase difference of ear motions on direction detection performance. In the upper panels, blue line indicates the left ear’s orbit given by 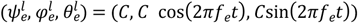, and orange line does the right ear’s orbit 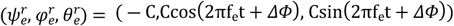, respectively. Black straight lines connect simultaneous points of the left and right ears’ orbits with the phase difference *ΔΦ*. In particular, the ear motion with *ΔΦ* = 180° is 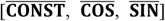 and the ear motion with *ΔΦ* = 0° is 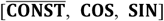. For motions with *ΔΦ* between 45° and 180°, good-quality direction detection performance is achieved.

## Discussion

In this study, we developed a theoretical model in which only certain ear motions consisting of three-axis rotations accomplished 3D direction detection accurately (Figs 8 and 10) and robustly (Fig 9). In the real world, bats intentionally employ rapid ear motions for 3D localization, despite the high energy costs, suggesting that they provide significant benefits in the process of echolocation.

Previous mathematical [27, 28] and practical demonstrations [29] have shown that ear motions can be useful under certain motion patterns. In contrast, our study has considered the theoretical basis for these ear motions by evaluating exhaustive motion patterns. Thus, this is the first article to investigate the underlying theory behind the ear motion strategies of bats. The results of general case analyses (Fig 10) show that three-axis rotations are necessary for 3D direction detection (i.e., those not including the pairing name **0**).

In particular, the pitch angle functions 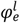 and 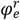 must retain a different phase, as shown in Fig 10. Such antiphase control of pitch motions has been observed in bats [19], and so our theory strongly supports the inevitability of pitch control in actual bat behavior. Our analyses indicate that the same antiphase control restriction exists in the yaw angle functions 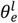 and 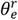, but the roll angle functions 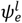 and 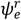 have no such restriction. These differences might be caused by the fact that pitch and yaw angles determine the central direction of the directivity pattern, while the roll angle determines the rotation around the direction axis. Thus, our investigations provide not only theoretical support for bats’ behavior, but also a new interpretation for roll–pitch–yaw controls. Such a cross-insights between theoretical and behavioral investigations reaches the core of active listening behaviors.

Ear motion patterns which give accurate and robust direction detection were only found in five of the 36 motion patterns analyzed in this study, as shown in Figs 8 and 9. This suggests that bats select a motion pattern from these five patterns. Thus far, we have neglected the physiological properties of bats in our analyses. It is plausible to assume that the motions of the left and right ears are mirror symmetric with respect to the surgical plane. If so, the following equations should hold:

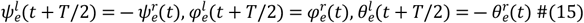

Only one of the five high-performance patterns satisfies the above equations, namely 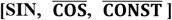. Therefore, we speculate that actual bats adopt ear motions that are close to this pattern (see the animation in **S1 Video**).

We have not only identified a wide array of patterns of appropriate ear motions (Fig 10), but have also provided simple discrimination conditions using orbits in the roll– pitch–yaw space. Our graph-based evaluation method is also useful for ethological investigation, because the graphs can be drawn using actual measurement data. Moreover, the hearing directivity pattern is also able to approximate from actual measurement data. Thus, we provide not only theoretical findings, but also an extendable framework of theoretical analysis for ethological research.

In our study, only one directionality of the ear and limited numbers of ear motions were investigated; thus, it is true that our theory is not perfect. However, the significance of this study lies in showing that simple shaped hearing directionality and well-selected uncomplicated ear motions are sufficient to achieve precise and robust direction detection. In addition, we proposed an index (degree of injection) that can judge whether the well-behaved inverse map is constructible or not using only the original map, without requiring the construction of an inverse map. Thus, we expect it to be useful for general-purpose evaluation systems in sensing fields.

## Acknowledgments

We are grateful to Toshihira Mishima for useful discussions and for providing useful insights that will serve as seeds for my research. We thank Stuart Jenkinson, PhD, from Edanz (https://jp.edanz.com/ac) for editing a draft of this manuscript.

## Supporting information

**S1 Video**. *Example movie for appropriate motion of left and right ears*.

**S1 Text**. *Echo amplitude representation procedure with four omni-directional microphones*.

**S2 Text**. *Expression of attitude of the directional ear*.

## Notes

### Competing Interest Statement

The authors have declared no competing interest.

## References

1. Griffin DR. Listening in the dark: the acoustic orientation of bats and men. 1958.

2. Fujioka E, Aihara I, Sumiya M, Aihara K, Hiryu S. Echolocating bats use future-target information for optimal foraging. Proceedings of the National Academy of Sciences. 2016:201515091.

3. Ghose K, Horiuchi TK, Krishnaprasad PS, Moss CF. Echolocating bats use a nearly time-optimal strategy to intercept prey. PLoS Biol. 2006;4(5):e108. doi: 10.1371/journal.pbio.0040108. PubMed PMID: 16605303; PubMed Central PMCID: PMC1436025.

4. Hase K, Kadoya Y, Maitani Y, Miyamoto T, Kobayasi KI, Hiryu S. Bats enhance their call identities to solve the cocktail party problem. Communications biology. 2018;1(1):1–8.

5. Ulanovsky N, Fenton MB, Tsoar A, Korine C. Dynamics of jamming avoidance in echolocating bats. Proceedings of the Royal Society of London Series B: Biological Sciences. 2004;271(1547):1467–75.

6. Carr CE, Konishi M. Axonal delay lines for time measurement in the owl’s brainstem. Proceedings of the National Academy of Sciences. 1988;85(21):8311–5.

7. Suga N. Cortical computational maps for auditory imaging. Neural networks. 1990;3(1):3–21.

8. Park TJ, Grothe B, Pollak GD, Schuller G, Koch U. Neural delays shape selectivity to interaural intensity differences in the lateral superior olive. Journal of Neuroscience. 1996;16(20):6554–66.

9. Joris PX, Yin T. Envelope coding in the lateral superior olive. I. Sensitivity to interaural time differences. Journal of neurophysiology. 1995;73(3):1043–62.

10. Park T, Monsivais P, Pollak G. Processing of interaural intensity differences in the LSO: role of interaural threshold differences. Journal of Neurophysiology. 1997;77(6):2863–78.

11. Goldberg JM, Brown PB. Functional organization of the dog superior olivary complex: an anatomical and electrophysiological study. Journal of neurophysiology. 1968;31(4):639–56.

12. Tsuchitani C, Boudreau JC. Single unit analysis of cat superior olive S segment with tonal stimuli. Journal of Neurophysiology. 1966;29(4):684–97.

13. Kulesza Jr RJ. Cytoarchitecture of the human superior olivary complex: medial and lateral superior olive. Hearing research. 2007;225(1-2):80–90.

14. Irving R, Harrison J. The superior olivary complex and audition: a comparative study. Journal of Comparative Neurology. 1967;130(1):77–86.

15. Aytekin M, Grassi E, Sahota M, Moss CF. The bat head-related transfer function reveals binaural cues for sound localization in azimuth and elevation. The Journal of the Acoustical Society of America. 2004;116(6):3594–605.

16. Obrist MK, Fenton MB, Eger JL, Schlegel PA. What ears do for bats: a comparative study of pinna sound pressure transformation in Chiroptera. Journal of Experimental Biology. 1993;180(1):119–52.

17. Firzlaff U, Schuller G. Directionality of hearing in two CF/FM bats, Pteronotus parnellii and Rhinolophus rouxi. Hearing research. 2004;197(1-2):74–86.

18. Jen PH-S, Chen D. Directionality of sound pressure transformation at the pinna of echolocating bats. Hearing research. 1988;34(2):101–17.

19. Griffin D, Dunning D, Cahlander D, Webster F. Correlated orientation sounds and ear movements of horseshoe bats. Nature. 1962;196(4860):1185–6.

20. Yin X, Müller R. Fast-moving bat ears create informative Doppler shifts. Proceedings of the National Academy of Sciences. 2019;116(25):12270–4.

21. Qiu P, Müller R. Variability in the rigid pinna motions of hipposiderid bats and their impact on sensory information encoding. The Journal of the Acoustical Society of America. 2020;147(1):469–79.

22. Pye J, Roberts L. Ear movements in a hipposiderid bat. Nature. 1970;225(5229):285–6.

23. Simmons JA. The resolution of target range by echolocating bats. The Journal of the Acoustical Society of America. 1973;54(1):157–73.

24. Mantani S, Hiryu S, Fujioka E, Matsuta N, Riquimaroux H, Watanabe Y. Echolocation behavior of the Japanese horseshoe bat in pursuit of fluttering prey. J Comp Physiol A Neuroethol Sens Neural Behav Physiol. 2012;198(10):741–51. doi: 10.1007/s00359-012-0744-z. PubMed PMID: 22777677.

25. Schnitzler H-U, Denzinger A. Auditory fovea and Doppler shift compensation: adaptations for flutter detection in echolocating bats using CF-FM signals. Journal of Comparative Physiology A. 2011;197(5):541–59.

26. Hiryu S, Shiori Y, Hosokawa T, Riquimaroux H, Watanabe Y. On-board telemetry of emitted sounds from free-flying bats: compensation for velocity and distance stabilizes echo frequency and amplitude. Journal of Comparative Physiology A. 2008;194(9):841–51.

27. Vanderelst D, Reijniers J, Steckel J, Peremans H. Information generated by the moving pinnae of Rhinolophus rouxi: tuning of the morphology at different harmonics. PloS one. 2011;6(6):e20627.

28. Vanderelst D, Holderied MW, Peremans H. Sensorimotor model of obstacle avoidance in echolocating bats. PLoS computational biology. 2015;11(10):e1004484.

29. Walker V, Peremans H, Hallam J. One tone, two ears, three dimensions: A robotic investigation of pinnae movements used by rhinolophid and hipposiderid bats. The Journal of the Acoustical Society of America. 1998;104(1):569–79.

30. Yamada Y, Mibe Y, Yamamoto Y, Ito K, Heim O, Hiryu S. Modulation of acoustic navigation behaviour by spatial learning in the echolocating bat Rhinolophus ferrumequinum nippon. Scientific reports. 2020;10(1):1–15.

31. Gao L, Balakrishnan S, He W, Yan Z, Müller R. Ear deformations give bats a physical mechanism for fast adaptation of ultrasonic beam patterns. Physical review letters. 2011;107(21):214301.

